# Identification of novel bone-resorption markers by osteoclasts *in vitro* and *in vivo*

**DOI:** 10.1101/2024.05.10.593489

**Authors:** BP Norman, JP Dillon, SM Alkharabsheh, PJM Wilson, AS Davison, E Chapman, J Baker, S Coyle, C Probert, LR Ranganath, JA Gallagher

## Abstract

Bone resorption involves dissolution of mineral and enzymatic degradation of bone matrix. The primary enzyme is cathepsin K but other proteases including matrix metalloproteinases are involved. Some cleavage products of cathepsin K have been partially identified, including crossed-linked telopeptides of type I collagen. However, the pathway of type I bone collagen degradation has not been fully elucidated. The aim of this study was to comprehensively characterise the entire complement of bone breakdown products resulting from osteoclast action under controlled conditions *in vitro*. Complete characterisation of these breakdown products will advance understanding of osteoclast biology and has the potential to reveal new biomarkers of bone resorption.

We analysed extracellular media from osteoclasts cultured on dentine substrates, using untargeted liquid chromatography mass spectrometry. We discovered 22 breakdown products resulting from osteoclastic action. These products were peptide fragment sequences that mapped to various collagen proteins present in bone and dentine matrix. Nine peptide fragments mapped exclusively to collagen I alpha-1 chain (COL1A1), the most abundant protein in bone. Analysis of the reported cleavage sites in the COL1A1 protein sequence indicated 7/9 COL1A1-specific fragments not explained by known proteolytic events. We subsequently showed that 14 of the fragment products were present in human serum and/or urine from metabolomic datasets obtained from patients with the inherited metabolic disease alkaptonuria (serum) and lung cancer (urine). Two products were at higher concentration (P <0.05, fold change >2) in urine from patients with bone metastasis (29/112) from the lung cancer cohort.

The range of collagen peptide fragments we discovered as a direct result of osteoclast activity indicates a complexity of bone resorption pathways not previously known. Monitoring the concentrations of these novel bone markers in biofluids has the potential to capture multiple pathways of bone resorption activity beyond the existing assays based on Cathepsin K.

**Lay summary:** Breakdown of bone tissue is performed by specialised bone cells called osteoclasts in a process known as bone resorption. Knowledge of specific molecules produced from osteoclasts acting on bone is important for a) understanding bone resorption in health and disease, and b) clinical tests of bone resorption from measurement of these breakdown products. Here we aimed to characterise the entire complement of bone breakdown products resulting from osteoclast action under controlled conditions in the laboratory. We found a total of 22 breakdown products produced from osteoclasts cultured on wafers of dentine, a tissue with almost identical composition to bone. Analysis of the structures of these products revealed fragments of varying size produced from digestion of specific proteins present in bone. We then showed that 14/22 bone protein fragments observed in our cell culture experiments were also present in human blood and/or urine. In our analysis of urine from patients with lung cancer, we found that two of the protein fragments we identified were higher in a subset (29/112) of patients with bone metastasis. Our findings provide new insights into the mechanisms of bone resorption and show the potential for monitoring the range of urine bone protein fragments in bone disorders.

## Introduction

Osteoclasts are the principal cells that break down bone. Osteoclast action is fundamental to bone homeostasis, with an essential role in maintenance, repair and remodelling of bone tissue. Osteoclasts resorb bone through producing various proteolytic enzymes and secreting hydrogen ions into the sealed extracellular compartment between the osteoclast and bone tissue matrix where resorption occurs.^1^ Bone resorption results in the breakdown of bone matrix proteins. The predominant protein in bone is type I collagen, which constitutes 90% of the organic component of bone matrix. Self-assembly of the two type I collagen chains, the alpha-1 (I) and alpha-2 (I) molecules, results in a protein structure comprising a triple helix with a coiled conformation.^2^

Specific degradation products of type I collagen produced during bone resorption by osteoclasts are established markers of bone turnover. Following release of these products from the tissue matrix, they enter the circulation and are then excreted in urine. Quantification of type I collagen breakdown products in biofluid compartments provides a non-invasive readout of bone resorption activity. This is the basis of several biochemical assays employed routinely in clinical management of bone disorders. The existing biochemical bone resorption markers produced by the enzyme Cathepsin K do not represent the entirety of bone breakdown pathways by osteoclasts. Multiple studies have shown that even in healthy subjects, human serum and urine contain a range of peptide products derived from collagens, particularly collagen I alpha-1 chain (COL1A1).^3–8^ The variety of urine collagen fragments indicates that the protease cleavage sites in type I collagen I are extensive and not limited to the well-characterised Cathepsin K cleavage sites.^5^ It is likely that some type I collagen fragments are products of other proteolytic enzymes. For example, a group of non-cysteine protease enzymes expressed in bone and known to act on collagen is the matrix metalloproteinases (MMPs).^9^

A recent study demonstrated the potential of an approach based on proteolysis of bone under controlled conditions in the laboratory combined with high-resolution mass spectrometry (MS) for characterising the global breakdown pathways of collagens.^10^ Digestion of human type I collagen with purified cathepsin K resulted in the production of more than one-hundred peptide products originating from each of the COL1A1 and COL1A2 proteins. These products included peptides from known cathepsin K cleavage sites in addition to an array of previously uncharacterised peptides. This indicates new cleavage sites not limited to the cross-linking telopeptide domains but also including the helical portion of these proteins. The degree to which these products reflect osteoclastic resorption as it occurs *in vivo* or if they are present in human blood or urine is not known.

To provide a more physiologically representative view of osteoclast resorption by proteases not limited to cathepsin K, we studied the resorption products generated by activated osteoclasts cultured on dentine substrates, a matrix composition almost identical to bone. We then established the extent to which these identified breakdown products were observed in human serum and urine. Understanding the complexity of bone resorption pathways is fundamental knowledge in basic bone biology and has potential to inform new approaches for monitoring and treatment of disorders associated with dysregulation of bone remodelling.

## Methods

### Chemicals and reagents

Human osteoclast precursors, culture medium and medium supplements were from Lonza (Slough, UK). Base medium was OCP™ basal medium. Supplements and growth factors were from the OCP™ growth medium SingleQuots™ kit. Final growth medium composition was 10% fetal bovine serum, 2 mM L-glutamine, 100 units/mL penicillin/streptomycin, 33 ng/mL macrophage colony-stimulating factor (M-CSF) and 66 ng/mL receptor activator of nuclear factor kappa-beta ligand (RANKL).

For liquid chromatography mass spectrometry (LC/MS) analysis, deionised water was purified in-house by DIRECT-Q 3UV water purification system (Millipore, Watford, UK). Methanol, isopropanol (Sigma–Aldrich, Poole, UK), formic acid (Biosolve, Valkenswaard, Netherlands) and ammonium formate (Fisher Scientific, Schwerte, Germany) were LC/MS grade. Reference mass correction solution was prepared in 95:5 methanol:water containing 5 mmol/L purine (CAS No. 120-73-0) and 2.5 mmol/L hexakis(1H, 1H, 3H-tetrafluoropropoxy)phosphazine (HP-0921, CAS No. 58943-98-9) (Agilent, Cheadle, UK).

### In vitro dentine resorption assay

Dentine disks were prepared from hippo and walrus tusks. Disks were placed into wells of a 96 well plate. Human osteoclast precursor cells were cultured and differentiated into osteoclasts according to manufacturer guidelines. Osteoclast precursors were seeded into 96-well culture plates onto hippo dentine, walrus dentine or plastic at a density of 10,000 precursor cells per well in 100 µL growth medium containing M-CSF and RANKL for osteoclast generation. At day 7 and 14 of culture, 50 µL (50%) and 100 µL (100%) of medium, respectively, from each well was replaced with fresh growth medium. After 19 days, culture medium was harvested and stored at -80 °C until analysis. Dentine wafers were fixed in phosphate buffered saline containing 10% formalin and stained with toluidine blue to quantify resorption by reflected light microscopy.^11^ Non-resorptive control conditions were osteoclast precursors cultured onto plastic discs (+RANKL / -dentine; N = 6) or without RANKL (-RANKL / +dentine; N = 6, 3 walrus disks, 3 dentine disks), or onto plastic and without RANKL (-RANKL / -dentine; N = 3). Additional +RANKL / +dentine cultures were treated with 10 fM – 100 µM zoledronic acid (6 concentrations, N = 3 per concentration) at day 2, to examine potential effects of anti-resorptive treatment.

### LC/MS analysis

Extracellular media samples were thawed at room temperature then diluted 1:3 media: deionised water (v/v) in individual 1.5 mL microcentrifuge tubes. Sample tubes were vortexed for 10 sec then centrifuged at 1500× *g* for 5 min. 50 µL of supernatant was transferred to a 150 µL 96-well plate (Agilent, Cheadle, UK) for analysis. Pooled samples were created for quality control (QC) purposes for each treatment condition in addition to an overall QC by pooling equal volumes of individual samples. Pooled samples were prepared in an identical way to individual samples.

Culture media analysis was performed using a published liquid chromatography quadrupole time-of-flight mass spectrometry (LC-QTOF-MS) acquisition method, which employed a 1290 Infinity II HPLC coupled to a 6550 QTOF-MS equipped with dual AJS electrospray ionisation source (Agilent, Cheadle, UK).^12^ Sample injection volume was 10 µL. Reversed-phase LC was performed on an Atlantis dC18 column (3×100 mm, 3 μm, Waters, Manchester, UK) maintained at 60 °C. Mobile phase composition was (A) water and (B) methanol, both with 5 mmol/L ammonium formate and 0.1% formic acid. The elution gradient started at 5% B 0–1 min and increased linearly to 100% B by 12 min, held at 100% B until 14 min, then at 5% B for a further 5 min. MS data acquisition was performed in positive ionisation polarity with mass range 50–1700 in 2 GHz mode with acquisition rate at 3 spectra/second. A reference mass correction solution was continually infused at a flow rate of 0.5 mL/min via an external isocratic pump (Agilent, Cheadle, UK) for constant mass correction (see preparation of reference mass correction solution). Capillary and fragmentor voltages were 4000 V and 380 V, respectively. Desolvation gas temperature was 200 °C with flow rate at 15 L/min. Sheath gas temperature was 300 °C with flow rate at 12 L/min, and nebulizer pressure was 40 psi and nozzle voltage 1000 V. Reference ions monitored were: purine (*m/z* 121.0509) and HP-0921 (*m/z* 922.0098).

Generation of compound fragmentation spectra was obtained from MS2 analysis of media samples with accurate mass precursor ion targets detailed in Table 1; no more than six compound targets per injection. Multiple fixed collision energies were applied; 10, 20 and 40 eV. Acquisition rates were 6 spectra/sec in MS1 and 4 spectra/sec in MS2.

**Table 1.**
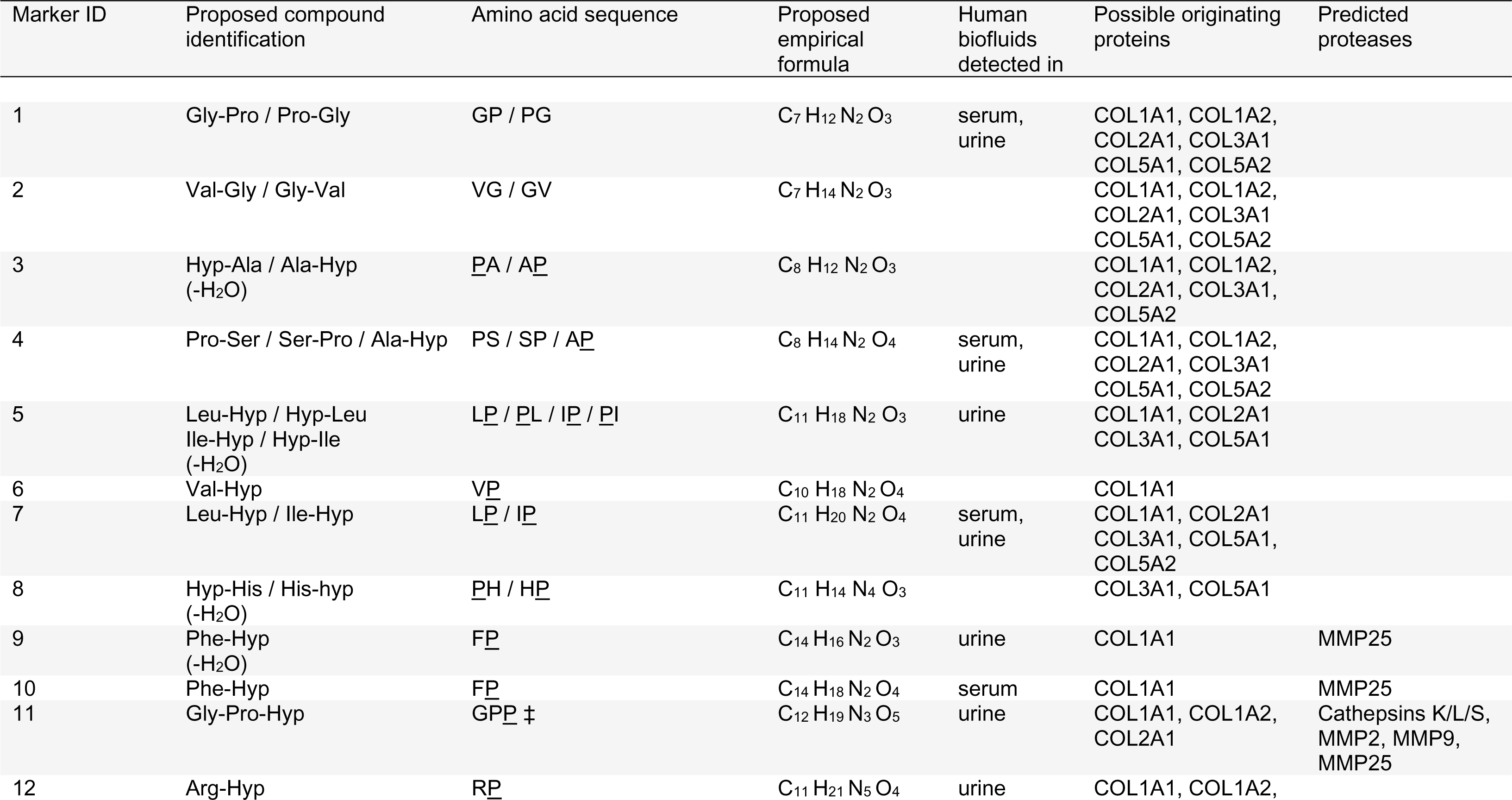
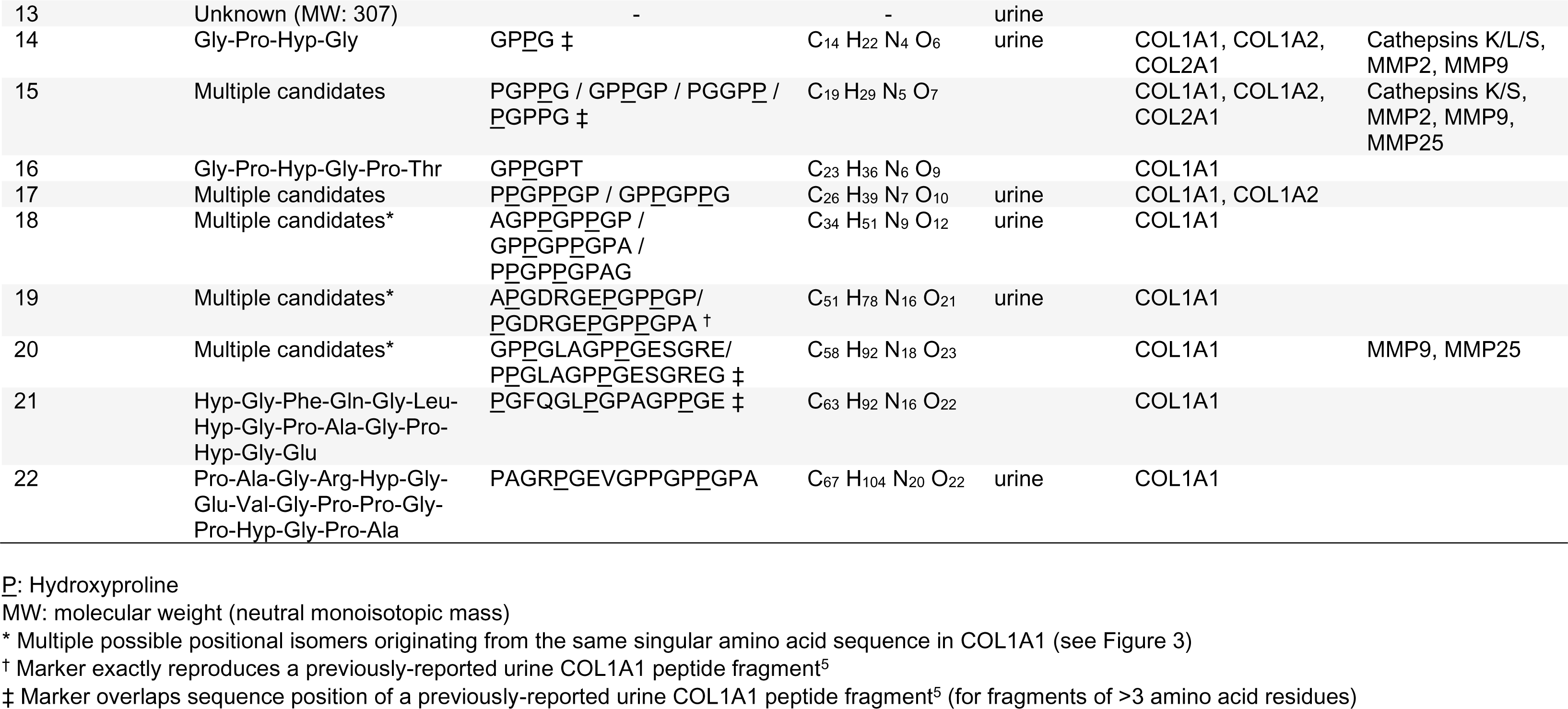
Summary of bone resorption products observed from LC/MS analysis. Proposed identifications (IDs) are based on mapping of observed peptide fragments against all possible oligopeptide products obtained from amino acid sequences for mature collagen proteins present in dentine. Predicted proteases are those proposed to produce the observed oligopeptides based on overlap between our markers and known protease cleavage sites in the COL1A1 protein; octapeptide cleavage sites obtained for cathepsin proteases and matrix metalloproteinases (MMPs) from the MEROPS^16^ database.

### Human metabolomic data

Human serum and urine metabolomic datasets were acquired under identical analytical conditions to those described above for *in vitro* experiment samples. These datasets are published and their associated raw data publicly available. Human urine data were obtained as part of a study investigating the metabolic changes that occur during the dying process in patients with incurable lung cancer (patients N = 112).^13^ Serial urine samples were collected from patients at various hospitals/hospices in North West England between 2016–2018 at various time points leading up to death (>12 weeks to <1 week before death); total samples N = 234. Among the 112 patients in the urine study, 29 patients had bone metastasis confirmed by CT, PET or bone scintigraphy; the other 83 patients had lung cancer but without known bone metastasis. Human serum data were obtained from the phase-3 trial of nitisinone treatment in patients with alkaptonuria, an inherited disorder of tyrosine metabolism (patients N = 103; 3 serial samples taken per patient across a 4-year period).^14^

### Data processing and statistical analysis

Raw LC/MS data were mined for compound features in Masshunter Profinder (build 10.00, Agilent, Cheadle, UK). Data from *in vitro* media samples were mined by an untargeted approach using the batch recursive (small molecules/peptides) feature extraction function. Untargeted feature extraction parameters: peak height >5000 counts, allowed ion species +H, +Na, +K and +NH_4_, alignment retention time (RT) tolerance ±0.3 min, alignment mass tolerance ±20 ppm, molecular feature extraction score >70 and Agile 2 peak integration method. Compounds not detected in all individual samples across at least one treatment condition were removed from the analysis.

Human metabolomic data were mined for the oligopeptide features identified from the *in vitro* experiment using the targeted feature extraction function in Profinder. Feature extraction parameters were accurate mass match window ±5 ppm and RT window ±0.3 min. Allowed ion species were +H, +Na, +K and +NH_4_. ‘Find by formula’ filters were: score >60 in at least 60% of samples from at least one sample group.

Extracted peak area intensity data were exported in .csv file format and imported into Mass Profiler Professional (MPP; build 15.1, Agilent, Cheadle, UK). In MPP, all data were log_2_ transformed and pareto scaled. Human urine data were normalised by urine creatinine concentration. Quality control of extracted features was performed on all datasets as previously described, based on reproducibility of signals across replicate injections of each pooled sample; features present in 100% of injections for at least one pooled sample, and with CV <25% across replicate injections were retained in the analysis.^12^ Statistical comparisons between *in vitro* experiment groups were performed in MPP by one-way ANOVA analyses with false-discovery rate (FDR) adjustment and post-hoc analysis by Tukey’s Honest Significant Different test. Volcano plot analysis combining fold-change (FC) and T-test analyses were performed on the urine dataset in MetaboAnalyst (version 6.0)^15^ to examine potential differences in urine compounds between samples from patients with *versus* without bone metastasis.

### Compound structure identification

Compound identification was performed using multiple chemical properties; accurate mass and MS2 fragmentation spectra of extracted features, in addition to the known amino acid sequences of the major extracellular matrix proteins present in dentine. Extracellular proteins mapped to were collagens 1, 2, 3 and 5; COL1A1, COL1A2, COL2A1, COL3A1, COL5A1 and COL5A2. Reference amino acid sequences for these proteins, with hydroxyproline and hydroxylysine post-translational modifications, were obtained from the NCBI protein database (National Library of Medicine; National Institutes of Health; U.S. Department of Health and Human Services) with the following accession numbers: NP_000079.2 (COL1A1), NP_000080.2 (COL1A2), NP_001835.3 (COL2A1), NP_000081.2 (COL3A1), NP_000084.3 (COL5A1), NP_000384.2 (COL5A2). The theoretical monoisotopic neutral accurate masses of all possible oligopeptide chain sequences 1-20 amino acids long for these proteins were mapped out in Microsoft Excel. Matching of calculated neutral masses of observed features was performed against these sequence mass databases generated for each candidate protein (mass match window ±5 ppm).

## Results

### Resorption of dentine discs in vitro

Extensive resorption of dentine disks was observed in cell cultures grown on dentine substrate with activation of osteoclast activity by RANKL supplementation in growth medium (+dentine / +RANKL). In these cultures, resorption pits covered over 60% of the dentine substrate surface area, as shown in Figure 1A. No difference was observed in the extent of resorption between hippo and walrus dentine. Resorption of dentine substrates was not observed in control cultures which had no dentine substrate and/or no RANKL supplemented in growth medium.

**Figure 1.**
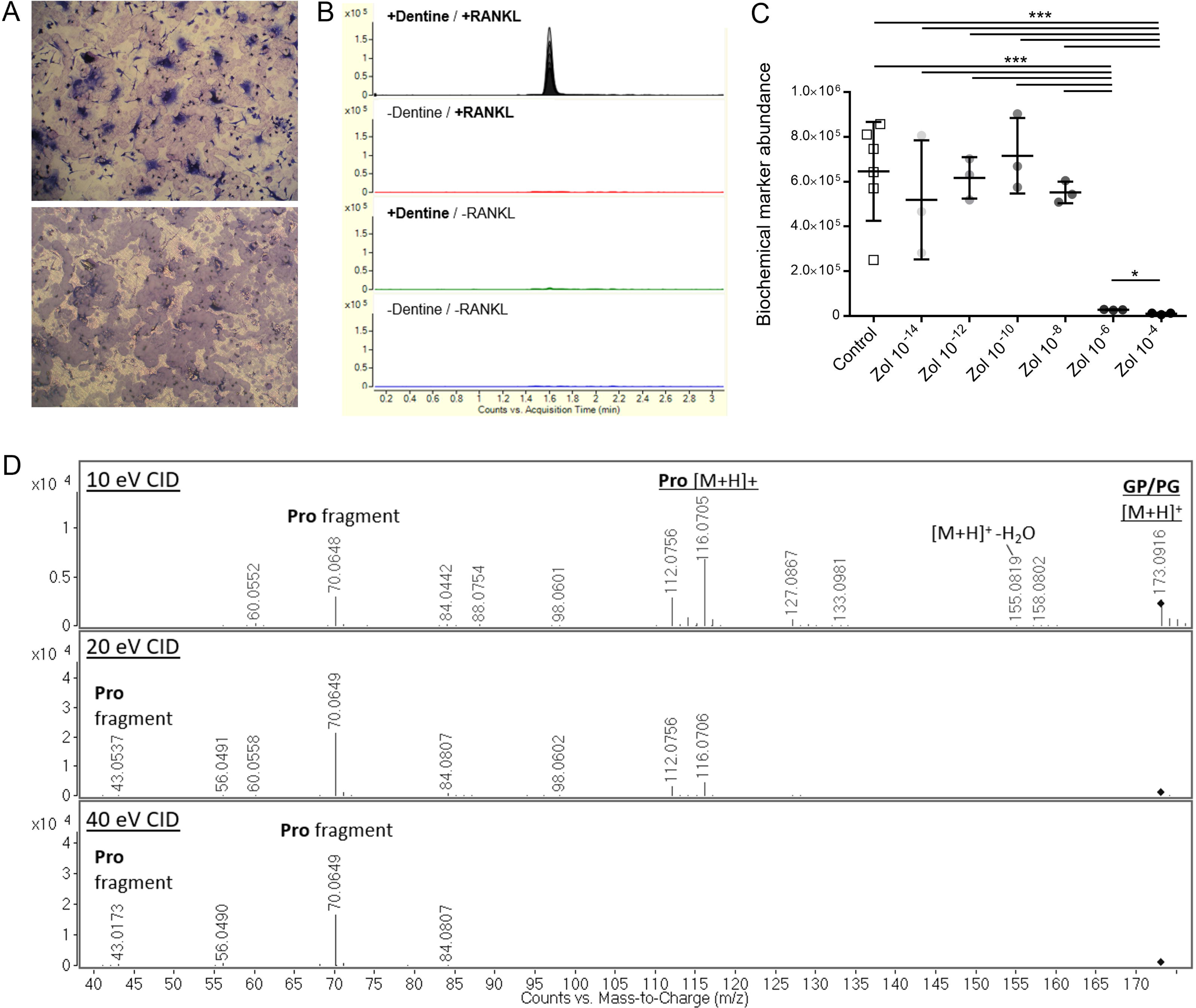
Observation of bone resorption in the dentine slice assay and identification of oligopeptide biochemical markers of bone matrix breakdown. **A**: Upper panel: Transmitted light photomicrograph showing osteoclasts resorbing dentine substrate. Lower panel: Reflected light photomicrograph showing resorption trails. **B**: Representative chromatogram of an oligopeptide bone resorption marker from extracellular media sampled from the dentine slice assay. Note the exclusive presence of signal in cultures of RANKL-activated osteoclast precursors grown on dentine but no signal detected in any of the control conditions. Traces from individual samples are overlaid within each treatment group. **C**: Dose-responsive inhibition of peptide fragment release into extracellular media with increasing concentrations of zoledronic acid treatment 10^-14^ M (10 fM) - 10^-4^ M (100 µM) in activated osteoclast cultures. Dot plot shows a representative example of the sharp decline in extracellular oligopeptide signal observed between 10^-8^ and 10^-6^ M zoledronic acid concentrations. * P <0.05; *** P <0.001. **D**: Example MS/MS fragmentation spectra for peptide fragment at multiple collision energies, showing the intact molecular [M+H]^+^ cation and MS peaks representing constituent substructures, including the individual amino acid proline. CID: collision-induced dissociation. The representative peptide fragment product shown in sections B-D is the dipeptide comprised of proline and glycine.

### Oligopeptide markers of in vitro bone resorption in extracellular media

Untargeted feature extraction of LC/MS data acquired from extracellular media resulted in 153 positively charged ion features across all samples. Quality control filtering resulted in 148 features present in all replicate injections for at least one pooled group sample; 147 of these features passed additional variability filtering, with CV <25% across replicate injections of each pooled sample. Thirty-one features had increased abundance (P <0.05, fold change >2) in resorption-positive +dentine / +RANKL osteoclast cultures compared with all control cultures. Twenty-two of these features were identified as potential resorption markers following additional QC checks on raw data for visualisation of feature integration quality and removal of duplicate extracted features (Table 1). Among these chemical entities, 21 compounds were exclusively present in +dentine / +RANKL cultures (Figure 1B), except for glycyl-prolyl-hydroxyproline, which was also detected in control cultures at lower abundance; raw peak area mean (±SD) = 364,326 (±73,573) in +dentine / +RANKL, mean (±SD) = 45,755 (±15,310) across control cultures (Figure S1).

The abundance of 21/22 resorption compounds showed a clear dose-responsive decrease with zoledronic acid treatment in +dentine / +RANKL cultures; P <0.0001. There was strong concordance between the profiles of these compounds, with a consistent sharp decline at zoledronic acid concentration ≥1 µM (Figure 1C). Post-hoc analysis confirmed this trend, with significant differences (P <0.01 or <0.001) between the 100 µM and 1 µM zoledronic acid groups compared with all lower zoledronic acid concentrations. Strong positive correlations were observed between marker abundance and the quantified resorption area of dentine discs across the zoledronic acid treatment groups (mean ±SD R^2^ = 0.72 ±0.11). Three compounds showed further decreases with zoledronic acid concentration between 1 µM and the highest dose of 100 µM (P <0.05); these were proline, glycine and hydroxyproline-containing oligopeptides with *m/z* 173.0925 (Pro-Gly or Gly-Pro), 343.1619 (Gly-Pro-Hyp-Gly) and 610.282 (Pro-Hyp-Gly-Pro-Hyp-Gly-Pro or Gly-Pro-Hyp-Gly-Pro-Hyp-Gly). Dose response to zoledronic acid treatment was not observed for the marker with *m/z* 251.1142 (His-Hyp or Hyp-His) despite not being detected in the -dentine / +RANKL, +dentine / -RANKL or -dentine / -RANKL controls (Figure S2).

### Chemical identification and mapping oligopeptide markers to matrix proteins

With the exception of one compound (compound #13, Table 1), all compounds mapped to one or more oligopeptide amino acid sequence present in the extracellular matrix proteins COL1A1, COL1A2, COL2A1, COL3A1, COL5A1 or COL5A2 (Table 1, Table S1). Eleven of the 22 compounds were identified as various di-peptides and mapped to multiple sequences across the proteins examined, except for three dipeptides that were unique to COL1A1. Nine of the 22 compounds mapped exclusively to the COL1A1 protein. These COL1A1-specific oligopeptides ranged from 2 (Val-Hyp and Phe-Hyp) to 17 amino acids. The five highest molecular weight products were all COL1A1 specific and mapped uniquely to single sequences within this protein (Figure 2). These COL1A1-specific compounds were doubly charged and contained a minimum of one Gly-Pro-Hyp sequence. Four compounds were identified as products of dipeptides minus the accurate mass of H_2_O. These molecules are unlikely to be technical artefacts caused by MS in-source fragmentation, as the chromatographic retention time was different to their derivative complete dipeptides where these were observed.

**Figure 2.**
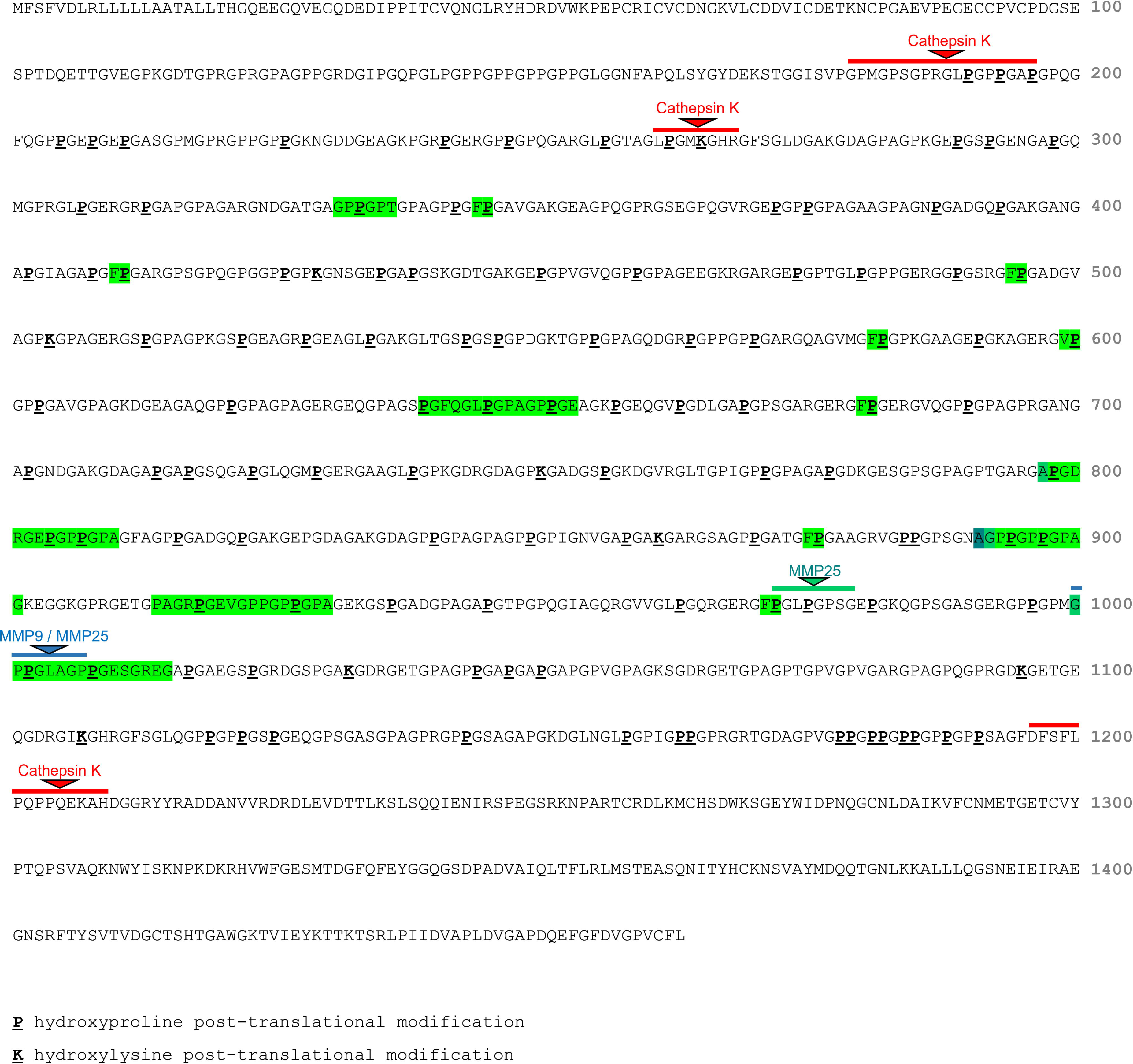
Mapping positions of peptide fragments, highlighted green, that mapped exclusively to the human COL1A1 protein sequence. Among these fragment sequences, 2/9 overlapped the positions of reported protease cleavage sites in COL1A1^16^; one cleavage site of MMP25 and another of both MMP9 and MMP25. The reported cleavage sites of Cathepsin K are shown for reference.

### Comparison of COL1A1 fragment peptide sequence locations to known COL1A1 cleavage sites

Known proteolytic cleavage sites in the COL1A1 protein were investigated to identify candidate collagenolytic enzymes with the potential to produce the COL1A1 peptide fragments discovered. Using the MEROPS database of proteolytic enzymes^16^, COL1A1 was selected as a substrate protein and all reported cleavage sites across all reported COL1A1-targeting peptidases were identified. These peptidases included cathepsins D, L, K and S and various MMPs. All COL1A1 cleavage sites reported from any form of evidence in this database were considered. All of our identified COL1A1 peptide fragments greater than three amino acid residues were considered in this analysis, with the exception of the di-peptides Phe-Hyp and Valyl-Hyp. These fragments, and particularly the longer sequences, were generally more specific to COL1A1.

Table 1 shows the predicted proteases associated with each peptide fragment based on overlap between their COL1A1 mapping positions and all known COL1A1 protease cleavage sites provided in MEROPS. The COL1A1 protein sequence positions of six of our peptide fragments overlapped with a known COL1A1 proteolytic cleavage site. These known cleavage sites are targeted by the collagenolytic enzymes Cathepsins K, L and S, MMP2, MMP9 and MMP25 (MMP25; previously known as MMP20). Figure 2 shows the COL1A1 sequence positions of the nine peptide fragments that mapped exclusively to COL1A1. Only 2/9 COL1A1-specific peptides overlapped with known cleavage sites. These were phenylalanine-hydroxyproline, which maps to multiple COL1A1 sequences, one of which overlaps an MMP25 cleavage site, and a 15-amino acid residue peptide fragment which overlapped with one cleavage site reported for both MMP9 and MMP25 (marker #20; see Table 1). None of our other observed peptide fragments greater than 5 amino acid residues overlapped a known COL1A1 cleavage site.

### Detection of novel oligopeptide markers in human serum and urine

Fourteen of the 22 peptide fragments we discovered from *in vitro* experiments were detected in human serum and/or urine; four fragments in serum, 13 fragments in urine (Table 1). The serum and urine metabolomic datasets were acquired under identical analytical conditions to those used in this study. Signals for these molecules were reproducible, with peak area CV <25% across replicate injections of various pooled samples across the analytical run (pooled urine profiles shown in Figure 3A). All serum compounds were also detected in urine, with the exception of Phe-Hyp, which was only reliably detected in serum. Gly-Pro-Hyp and an unidentified compound with *m/z* 308.122 (compound #13; Table 1) were more abundant (P <0.05, fold change >2) in urine from patients with bone metastasis among the lung cancer cohort; n = 29 patients with lung cancer and bone metastasis; n = 83 patients with lung cancer but without known bone metastasis (Figure 3B).

**Figure 3.**
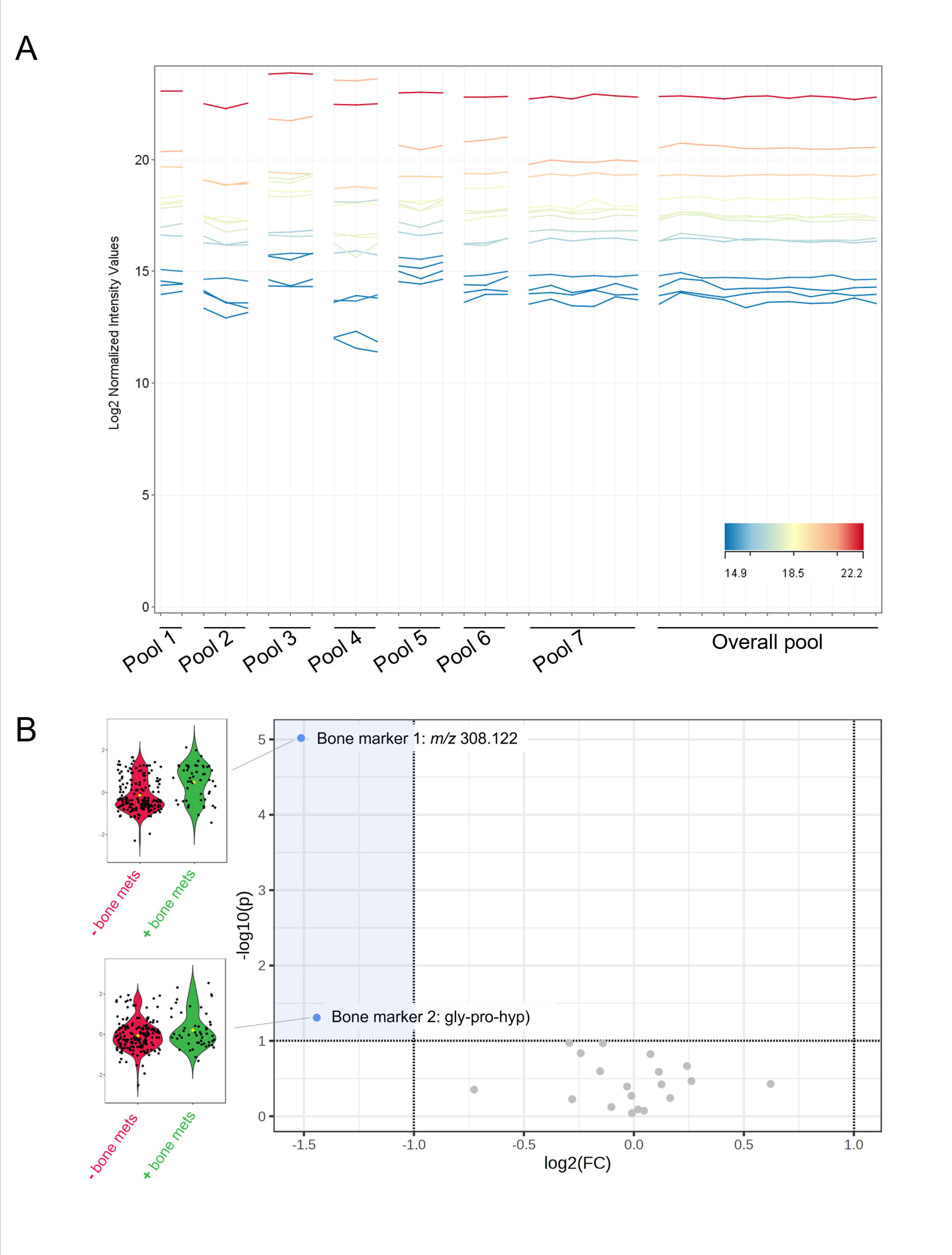
Oligopeptide products as potential urinary markers of bone resorption and metastatic bone disease in humans. **A**: Abundance profiles of 13 oligopeptide fragments reliably detected in human urine. Profiles are extracted peak areas of individual compounds across multiple injections of eight pooled samples representing each biological group (time point before death; <1 week to >12 weeks from death) from a published urine metabolomic study.^13^ Replicate injections of each pooled sample were randomly interspersed across the entire analytical run which comprised 297 injections. For all peptide fragments, CV was <25% across replicate injections for each pooled sample. **B**: Volcano plot highlighting the two oligopeptide compounds altered in urine from patients with bone metastasis among the lung cancer cohort (blue blots in shaded region indicating P <0.05 and fold change >2 between patients with bone metastasis *versus* controls). These compounds were Gly-Pro-Hyp and an unidentified compound with m/z 308.122 (compound #13, Table 1). All remaining urinary oligopeptides detected showed no significant difference (P >0.05 and fold change <2) between patient groups (grey dots). N = 29 patients with lung cancer and bone metastasis (mets), n = 83 patients with lung cancer but without known bone metastasis.^13^

## Discussion

Our data show culturing osteoclasts on dentine substrate provides a useful tool to model bone resorption as it occurs *in vivo*. Fourteen of the 22 bone breakdown peptide products released into culture medium and discovered under controlled conditions in this model system were subsequently detected in human serum and urine under identical LC/MS analytical conditions. Four of the 22 fragments were found in serum in patients with alkaptonuria, an inherited disorder of tyrosine metabolism with joint pathology as a main feature. Thirteen fragments were found in urine from patients with lung cancer, and 2/13 were higher in urine among a subgroup of these patients with bone metastasis.

Consistent with previous studies^3–8^, our observed bone peptide fragment products were rich in hydroxyproline residues and more frequently observed and at higher abundance in urine *versus* serum. The products spanned a wide range in molecular weight (174-1540 Da), from di-peptides to peptide fragments comprising 17 amino acid residues (Table 1). The lowest molecular weight products, mainly di- and tri-peptides, mapped to various sequences across the collagen proteins present in dentine, with the exception of the di-peptides Phe-Hyp and Val-Hyp which were specific to COL1A1. In total we identified 9/22 peptides with sequences mapping exclusively to COL1A1, the protein that comprises two of the three chains in the type I collagen triple helix and the most abundant protein in dentine and bone. Amongst the COL1A1 specific products, two overlapped with the sequences of previously reported urine COL1A1 fragments and one reproduced exactly another previously reported fragment.^5^ Three of the COL1A1-specific resorption products are not previously described. Six of the seven highest molecular weight products did not map to known COL1A1 protease cleavage sites, potentially indicating new COL1A1 proteolytic degradation pathways.

The peptide fragments we discovered indicate new cleavage sites in type I collagen. As shown in Figure 3, 7/9 fragment sequences identified as COL1A1-specific did not overlap the positions of cleavage sites reported in MEROPS^16^, one of the major protease databases. Instead, these peptide fragments mapped to more central positions within COL1A1 between the C- and N-terminal telopeptide sites at the extremities of the protein. Our observation adds to previous reports of type I collagen peptides in urine that do not seem to result from known proteolytic events.^5^ Compared with previous analyses of urine peptides, we observed more low molecular weight markers such as di- and tri-peptides. These smaller fragments were less specific, usually mapping to multiple sequences within COL1A1, including cathepsin K cleavage sites, in addition to sequences within the other collagen proteins we considered (Table 1, Figure S2). Observation of small collagen fragments indicates that our *in vitro* experiment procedure, involving analysis of extracellular media content after days of osteoclast activity, more represents end-stage resorption. Controlled digestion of type I collagen with cathepsin K in a previous study showed a chronology of proteolysis, with early cleavage events after one hour of digestion occurring preferentially at the telopeptide cross-linking domains at the extremities of the type I collagen molecule. With increasing digestion time, up to 16 hours, cleavage occurred across the breadth of collagen, including the central helical domain. Over this longer digestion period, more isolated fragments were observed including peptides of fewer than four amino acid residues.^10^ The extent to which this non-specific cleavage by cathepsin K over longer periods of digestion explains the smallest peptide fragments we observed in our study is not clear.

Our data support biofluid collagen peptide fragments as potential biomarkers in disorders affecting bone. Two of our urinary compounds showed increases in a subset of patients with bone metastasis (29/112) among our patient cohort with advanced lung cancer. Bone is a common site of metastases in most solid tumours including lung, breast, prostate and colorectal cancers. There is urgent need for non-invasive biochemical markers for early diagnosis and monitoring of bone cancer. An osteolytic phenotype is frequently induced by presence of bone tumours. Our finding of elevated urine bone markers is consistent with an extensive body of evidence showing type I collagen markers are higher in bone metastasis^17–19^. This increase in type 1 collagen markers is associated with stimulation of osteoclastogenesis through RANKL expression and increased cathepsin K activity.^20–22^ In our lung cancer cohort, we previously showed changes in a number of our bone markers in the last weeks of life.^13^ Two of the COL1A1-specific collagen peptides reported here progressively increased in urine over the course of time leading up to death, from greater than three months up to the last week before death; these were the dipeptide phenylalanyl-hydroxyproline and the highest molecular weight COL1A1 fragment observed here (marker #22; Table 1). Another unidentified bone marker (marker #13) decreased leading up to death. These changes provide major insights into the fundamental biological changes that occur at the end of life, implicating bone resorption pathways as part of a multifactorial process of dying. Numerous other studies show the clinical relevance of urine type I collagen peptide fragments in a variety of disorders including cardiovascular disease^6,23^, chronic kidney disease^7^, diabetes mellitus^3^, liver fibrosis^8^, and colorectal liver metastasis^24^. It is important to consider that even COL1A1-specific markers are not necessarily bone-specific *in vivo*, given the expression of type I collagen in most connective tissues and potential for urine collagen fragments to originate from the diet. However, Marx et al.^5^ provide evidence that the urine type I collagen peptides reported in their study on kidney transplant recipient patients reflect skeletal processes. First, all COL1A1 markers correlated with serum CTX-I, helping to rule out that the urine collagen fragments were produced in the kidney. Second, 13 COL1A1 fragments showed a consistent decrease after bone-targeted anti-resorptive treatment with bisphosphonates.

The extent of type I collagen fragments we observed highlights the complexity of bone resorption. The multitude of pathways of type I collagen degradation raises the possibility that different conditions of bone dysregulation may be associated with alterations in different resorption pathways. Monitoring an array of type I collagen fragments produced by different resorptive mechanisms and proteolytic enzymes in patients has potential to provide a more detailed readout of bone turnover and homeostasis than existing assays. There is potential value in monitoring different collagen fragments produced by specific MMPs. Our data, for example, found one COL1A1-specific fragment (marker #20, Table 1) to be a product of MMP9/MMP25 cleavage. The MMPs, a family of zinc-dependent proteolytic enzymes, have a major role in bone development and maintenance. MMPs not only act as extracellular proteases targeting matrix proteins such as collagens, but are also key mediators of cellular processes including proliferation, differentiation and motility in addition to angiogenesis, apoptosis and wound healing.^25^ Altered activity of specific MMPs is a major factor in the pathogenesis of various common diseases associated with bone dysregulation, including osteoarthritis, bone cancer and a range of conditions associated with osteolysis^26^. In diabetes mellitus, for example, lower levels of some urine COL1A1 fragment peptides are consistent with reduced activity of specific MMPs such as MMP2, MMP3 and MMP9^27,28^, possibly attributable to lower rate of type I collagen cleavage by these proteases.

Previous analyses of type I collagen peptides show considerable variation in the fragments observed. Many factors are likely to account for this variation, including the experimental conditions used in studies of bone or collagen digestion *in vitro*, in addition to instrumentation and analytical conditions and workflows employed in detection of peptides. The LC/MS technique used here is different to platforms used in other studies reporting collagen peptide fragments, which have frequently used capillary electrophoresis coupled to mass spectrometry (CE/MS).^4–8,23^ Our LC/MS approach has been employed extensively in small molecule analyses including metabolomic applications and is possibly more optimal for detection of low, *versus* high, molecular-weight peptides.^12^ This contrasts the proteomic approaches used in previous studies and shows the potential to detect low-medium molecular weight bone collagen fragments in serum and urine metabolomic datasets. The data mining approach in terms of initial molecular feature extraction on raw LC/MS data was untargeted, in contrast to other studies reporting collagen-derived peptides. Identification of molecules of interest (*i.e.* bone resorption products) was performed *post hoc*. In this way, the extraction of signals/peptide features did not depend on existing knowledge including the sequences of known peptides and data from existing peptide libraries. In other words, the profiles of ion features that were detected and extracted from the raw data, independent of their subsequent biochemical identifications, were included in the analysis. An advantage of this untargeted approach is that some peptide features remain incompletely characterised and are therefore not detectable using common proteomic data mining approaches which rely upon existing peptide libraries. For type I collagen, incompletely characterised features include the extent of cross-links, cross linking is known to occur at multiple telopeptide and helical regions, and the entire array of post-translational modifications including glycosylation.^10^

In conclusion, our data demonstrate the powerful approach of untargeted high-resolution MS analysis for characterisation of the range of type I collagen degradation products. The overlap between collagen fragments observed *in vitro* and subsequent detection in human serum and urine demonstrates the culturing of osteoclasts on bone collagen substrate is a physiologically relevant model system for studying bone resorption. The novel bone markers presented, some of which are COL1A1-specific, provide new insights into the pathways of collagen degradation and support the clinical potential of monitoring collagen peptide fragments in bone disorders.

## Acknowledgements

The authors thank Agilent Technologies UK Ltd for kindly providing access to instrumentation for analysis of *in vitro* experiment samples in this study, with special thanks to Dr Gordon Ross of Agilent Technologies for his assistance in these analyses.

## Ethical approval statement

Biology of dying urine metabolomic dataset: ethical approval was provided by North Wales (West) Research Ethics Committee (REC reference 15/WA/0464).

SONIA 2 trial serum metabolomic dataset: ethical approval for the study was obtained from each of the participant sites including the North West—Liverpool Central Research Ethics Committee (reference: 13/NW/0567), the NURCH Ethical Committee in Piešťany (reference number: 04196/0029/001/001) and the EC Ile De France II committee in Paris (reference number: 2013-08-08).

