## Supplemental for "Identification of novel bone-resorption markers by osteoclasts *in vitro* and *in vivo*"

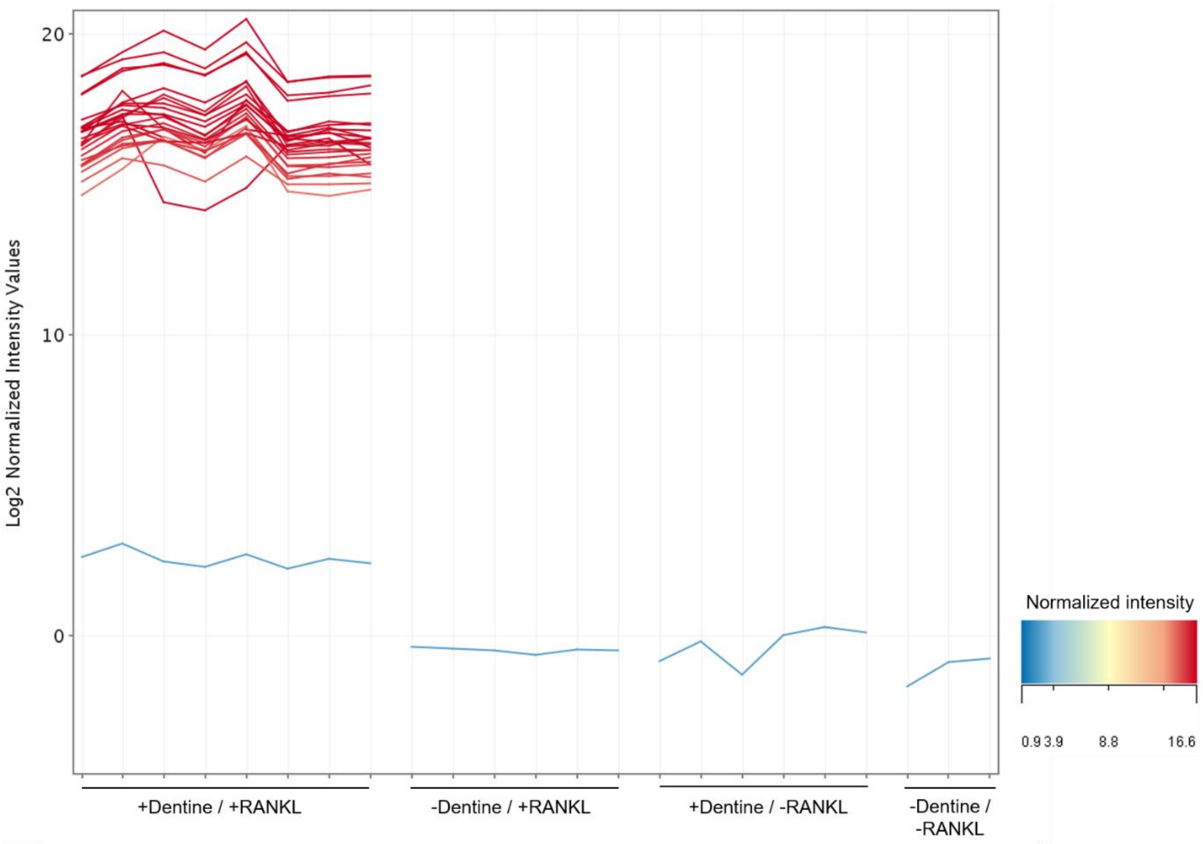


**Figure S1**. Profile plot showing extracellular media LC/MS peak area abundances of the 22 bone resorption products across the *in vitro* experiment conditions. Each line represents one compound. A clear cluster of compounds was observed representing products only detected in resorptive conditions (+dentine / +RANKL cultures); red profiles, top left portion of plot. The tripeptide Gly-Pro-Hyp (blue profile, lower portion of plot) was detected in all conditions, but with markedly greater abundance in +dentine / +RANKL compared with the three control groups.

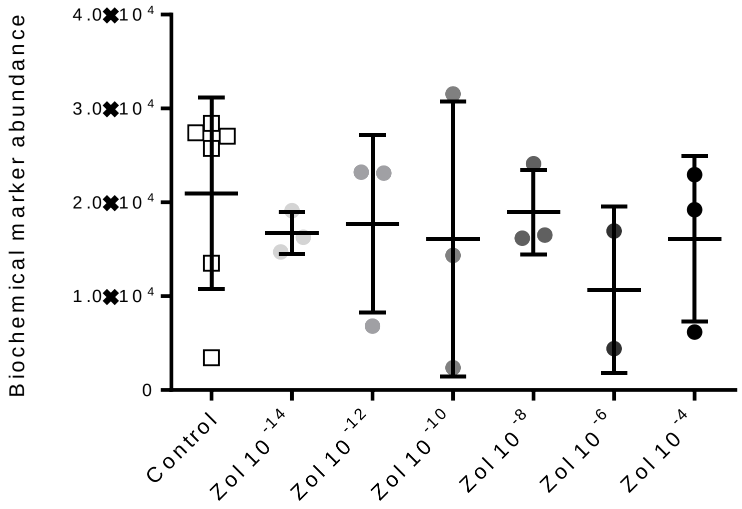


**Figure S2**. Peak area abundance values for hyp-his (or his-hyp); the only oligopeptide fragment observed in the dentine slice assay but without dose-responsive inhibition by zoledronic acid.

**Table S1.** Accurate mass and retention time data for bone breakdown products identified.

| Peptide | Measured *m/z* | Retention time (minutes) | Charge (*Z*) |
|---|---|---|---|
| GP / PG | 173.0925 | 1.6 | 1 |
| VG / GV | 175.1102 | 1.89 | 1 |
| PA / AP | 185.0926 | 2.58 | 1 |
| PS / SP / AP | 203.1032 | 1.36 | 1 |
| LP / PL / IP / PI | 227.1392 | 5.97 | 1 |
| VP | 231.1344 | 1.93 | 1 |
| LP / IP | 245.1501 | 3.29 | 1 |
| PH / HP | 251.1142 | 1.54 | 1 |
| FP | 261.1245 | 6.4 | 1 |
| FP | 279.1345 | 4.25 | 1 |
| GPP | 286.1431 | 1.74 | 1 |
| RP | 288.1674 | 1.27 | 1 |
| Unknown | 308.122 | 1.73 | 1 |
| GPPG | 343.1619 | 1.73 | 1 |
| PGPPG / GPPGP / PGGPP / PGPPG | 440.2132 | 3.78 | 1 |
| GPPGPT | 541.2603 | 3.76 | 1 |
| PPGPPGP / GPPGPPG | 610.282 | 3.78 | 1 |
| AGPPGPPGP / GPPGPPGPA / PPGPPGPAG | 389.6906 | 4.71 | 2 |
| APGDRGEPGPPGP/ PGDRGEPGPPGPA | 626.2847 | 4.44 | 2 |
| GPPGLAGPPGESGRE/ PPGLAGPPGESGREG | 705.3379 | 5.95 | 2 |
| PGFQGLPGPAGPPGE | 713.3364 | 5.6 | 2 |
| PAGRPGEVGPPGPPGPA | 771.3841 | 5.92 | 2 |
